# Modeling the kinetics of amyloid beta pores and long-term evolution of their Ca^2+^ toxicity

**DOI:** 10.1101/2022.05.02.490365

**Authors:** Syed Islamuddin Shah, Angelo Demuro, Ghanim Ullah

## Abstract

The formation of cation-permeable pores due to amyloid beta (Aβ) in the plasma membrane (PM) and membranes of intracellular organelles has been shown to cause disruption of Ca^2+^ homeostasis in Alzheimer’s disease, leading to neuronal malfunction and degeneration. However, detailed information about the formation, function, and time-evolution of Aβ pores are still lacking. Here we present a comprehensive analysis to show that the Ca^2+^ toxicity of PM pores formed by Aβ42 progressively increases over the duration of the experiments that last for more than an hour. This mainly results from a progressively increasing number of pores and activity of individual pores having higher open probability (P_o_) and Ca^2+^ conductance. To gain deeper insight into the function and evolution of Aβ42 pores and establish their role in Ca^2+^ toxicity beyond experimental times scales, we developed a data-driven computational model that not only mimics the kinetics of these pores at a given time but also tracks their time-evolution over many hours. Our model suggests that the rising toxicity of Aβ42 pores over time is a combined result of the large number of short-lived pores with low Ca^2+^ conductance and long-lived higher-conductance pores but fewer in number. Analysis and modeling of the voltage-dependent data show that Aβ42 pores are most active when the cell’s membrane potential is near resting values, making the cell at rest especially more prone to Ca^2+^ disruptions.

---

Alzheimer’s disease (AD) is the most common form of dementia inflicting a heavy socioeconomic toll [1, 2]. According to the “amyloid cascade hypothesis of AD” β-amyloid (Aβ) deposits drive formation of senile plaques and tau neurofibrillary tangles and thence neurodegeneration [3-6]. However, substantial new evidence implicates the soluble forms of Aβ oligomers, rather than the plaques, as the toxic specie in the pathogenesis of AD. One key mechanism through which soluble Aβ oligomers mediate their effects is by disrupting the integrity of cells plasma membrane, leading to uncontrolled flux of Ca^2+^ into the cells [7-18]. Various mechanisms have been proposed to underlie the increased permeability of plasma membrane to Ca^2+^, including interaction with several Ca^2+^ channels [13, 14, 16, 19-24]. However, studies on lipid bilayers [25-28] and *Xenopus* oocytes [17, 29, 30], which lack native plasma membrane Ca^2+^ channels, point to the formation of Ca^2+^-permeable pores by Aβ oligomers in the cell membrane as a major mechanism. Pore formation by Aβ has also been observed in HEK293 cells of neuronal origin [31]. By promoting uncontrolled influx across plasma membrane, these pores destabilize cell’s ionic homeostasis, particularly Ca^2+^, and make the basis for what is called the ion-channel hypothesis of AD [25, 32].

Significant progress has been made in establishing the gating properties of Aβ pores [7, 25, 33-38], their contribution to AD [39, 40], and their inhibition by various blockers, particularly zinc [41-45]. However, questions such as how the kinetic and Ca^2+^ toxicity of these pore evolve over longer time and how their function changes as the potential difference across cell’s membrane varies, remain unanswered. The importance of understanding the formation, functioning, and evolution of Aβ pores is further highlighted by the fact that oral application of the compound anle138b, which blocks the activity of conducting Aβ pores, before or after the onset of pathology, restores hippocampal synaptic and transcriptional plasticity as well as spatial memory in mouse model of AD [40].

In a seminal work, using high throughput and massively parallel optical patch clamp technique [46], we established that PM Aβ42 pores are highly dynamic [47], and leads to Ca^2+^ toxicity that significantly accelerates apoptosis [48]. Activity of thousands of these pores studied over tens of minutes shows that not only the number of Aβ40 pores increase over time, but the Ca^2+^ toxicity of existing pores continues to evolve due to increase in their open probability and Ca^2+^ permeability over time [47]. These advances have greatly improved our understanding of the functional properties of these pores, but they are limited to tens of minutes, and are unable to corroborate on how much exactly these pores contribute towards AD that spans over several decades. Thus, to establish the role of these pores in the disease progression, it is imperative to relate the progressive changes in their kinetics and permeability beyond experimental time scales [49]. Data-driven computational modeling is viable alternative to probe such experimentally inaccessible spatial and temporal timescales. Specifically, Markov chain modeling has proven to be an important tool in predicting the long-term behavior of biological systems employing experimental observations [50-52].

In this paper, we use the fluorescence time traces from thousands of Aβ42 pores recorded at successive times (see Materials and Methods section) after application of soluble Aβ42 oligomers to Xenopus Laevis oocytes, to develop the first model for the gating and time-evolution of Aβ42 pores. We have employed a pipeline of indigenously developed computational tools to extract (CellSpecks) [53] and idealize (TraceSpecks) [54] these time traces to identify various sub-conductance levels (SCLs) in which these pores can gate at a given time. These idealized traces representing the gating state of a pore as a function of time are then used to determine model parameters in terms of state-to-state transition rates using open source software QUB (Quantify Unknown Biophysics) [55]. Next, we use non-linear Hill like functions to fit these transition rates as functions of time. We have also probed the effect of hyperpolarizing voltage on the activity of these pores. By employing theory of aggregated binary reversible Markov chains [50, 56-58] we have used these fits to predict the effect of Aβ42 pores on Ca^2+^ homeostasis at any time (days and weeks) after cell’s exposure to the oligomers, allowing to probe their possible pathological connections towards AD progression.

## Materials and Methods

Aβ42 pores’ activity was imaged in the plasma membrane of *Xenopous laevis* oocytes at the single channel level using TIRFM and the data were saved in multi-frame MetaMorph stack files. Stack files were then processed and analyzed using our in-house software called CellSpecks for automatic detection of pores location and extracting gating properties [59]. Further statistical analysis was done in Matlab. Full details of our experimental approach are reported in [36] and are summarized below.

### Preparation and characterization of soluble Aβ42 oligomers

Solution of 0.5 mg of human recombinant Aβ42 peptide in 20 μl of freshly prepared DMSO were separately diluted with 480 μl of double-distilled water in a siliconized Eppendorf tube. After 10-min sonication, samples were incubated at room temperature for 10 min and then centrifuged for 15 min at 14,000 g. The supernatant fraction was transferred to a new siliconized tube and stirred at 500 rpm using a Teflon coated microstir bar for 8–48 h at room temperature. Aliquots were taken at intervals and were assayed by pipette application to voltage-clamped oocytes at a final bath concentration of 1 μg/ml. Membrane currents were recorded 20 to 25 min after application in response to hyperpolarization from 0 to -100 mV. Aβ42 preparations incubated for <6 h evoked little or no current, indicating that the monomeric peptide was ineffective at inducing membrane Ca^2+^ permeability. Preparations that evoked currents *≥* 1 μA at membrane potential of -80 mV were used immediately for TIRFM or were stored at -20°C before use [7].

### Oocyte preparation and electrophysiology

Experiments were performed on defolliculated stage VI oocytes. To visualize Ca^2+^ fluxes, oocytes were injected with Ca^2+^ sensitive dye fluo-4 dextran 1 h before imaging (molecular mass of 10 kD and Ca^2+^ affinity of 3 μM) to a final intracellular concentration of 40 μM. Before TIRF experiments, oocytes were placed in a hypertonic solution (200 mM K aspartate, 20 mM KCl, 1 mM MgCl2, 10 mM EGTA, and 10 mM Hepes, pH 7.2) at 4°C to shrink them so that the vitelline envelope could be manually torn apart and removed using fine forceps. Oocytes were then placed animal hemisphere down in a chamber whose bottom is formed by a fresh ethanol washed microscope cover glass (type-545-M; Thermo Fisher Scientific) and were bathed in Ringer’s solution (110 mM NaCl, 1.8 mM CaCl_2_, 2 mM KCl, and 5 mM Hepes, pH 7.2) at room temperature (23°C) continually exchanged at a rate of 0.5 ml/min by a gravity-fed superfusion system. The membrane potential was clamped at a holding potential of 0 mV using a two-electrode voltage clamp (Gene Clamp 500; Molecular Devices) and was stepped to more negative potentials (−100 mV unless otherwise indicated) when imaging Ca^2+^ flux through Aβs pores to increase the driving force for Ca^2+^ entry in to the cytosol. Solutions containing Aβs oligomers were delivered from a glass pipette with a tip diameter of 25-30 μm positioned near the membrane footprint of the oocyte membrane on the cover glass.

### TIRF microscopy and image acquisition

Imaging was accomplished by using a custom-built TIRFM system based around a microscope (IX71; Olympus) equipped with a 60× TIRF microscopy objective (1.45 NA; Olympus) [60]. Fluorescence excited by a 488-nm laser was imaged using an electron-multiplied charge-coupled device camera (Cascade 128+; Roper Scientific) at full resolution (up to 128 × 128 pixel; 1 pixel = 0.33 μm at the specimen) at a rate of 500 s^-1^. Image data were acquired using the MetaMorph software package (Universal Imaging) and were black-level corrected by subtracting the camera offset. The maximum observed fluorescence signals were small (maximum (F-Fo)/Fo < 2.0, where F is the fluorescence intensity at any time t and Fo is the baseline fluorescence intensity without any channel activity) in comparison to the full dynamic range of fluo-4 ((F-Fo)/Fo >30 in saturating Ca^2+^) and are thus expected to be linearly proportional to Ca^2+^ flux.

After application of Aβ42 solution, we performed several experiments to record the activity of these pores (and save them as Metamorph movie stacks) as a function of time and membrane potential [47, 48]. To evaluate the time dependent evolution of the activity of these pores, movie stacks from the same membrane patch were recorded at different times after application of Aβ42 solution with the first recording starting at 25 minutes (and lasting for 30 seconds). The subsequent stacks were recorded at 5 minute intervals with the last recording at 80 minutes after Aβ42 application.

### Processing movie stacks

Movie data on the activity of Ca^2+^ conducting Aβ42 pores often contains of several thousands of these pores (Figure 1A) conducting in various SCLs with tens of thousands of events ranging from few to several milliseconds. We have used our in-house software package called, CellSpecks, to process these movie records [53]. CellSpecks is capable of identifying the raw time traces of these individual pores, their locations, open and close events along with their durations, mean open (τ_o_) and close (τ_c_) durations, P_o_, maximum flux (amplitude) during each opening event, and the highest Ca^2+^ flux (maximum amplitude) ever observed for a pore during a single stack. Details of how CellSpecks perform these operations and the algorithms it uses to process these movie stacks are described elsewhere [61].

**Figure 1.**
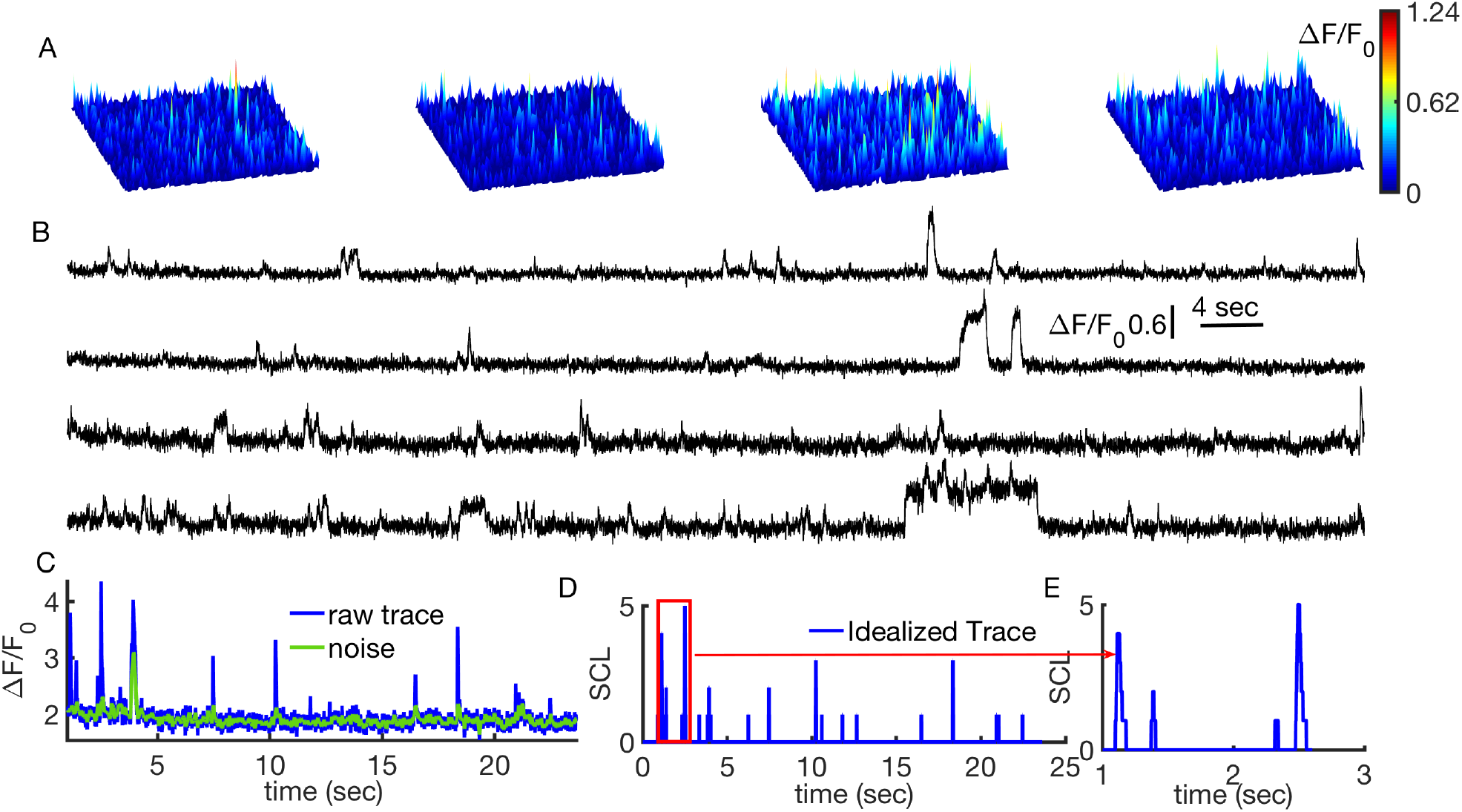
Processing raw movie stacks for the activity of Aβ42 pores. **A)** snapshots of the activity of Aβ42 pores from a typical movie stack recorded at a given time after cell’s exposure to Aβ42 oligomers (see Table 1). **B)** Representative raw time traces from 4 different pores. Each trace is retrieved from an area of ∼1 μm^2^. **C)** Raw time trace (blue) of an example Aβ42 pore as retrieved by CellSpecks. Green trace is the noise identified by TraceSpecks during idealization process of the raw (blue) trace. **D)** Resulting idealized trace, showing up to 5 SCLs of the pore in **(C). E)** A zoomed in version of the idealized trace showing the microscopic opening and closing of the pore in various SCLs.

Figure 1A shows example snapshots from a 25×25 μm^2^ patch of membrane over successive times produced by CellSpecks. Activity of each pore is represented by the peaks where bright colors represent pores with higher activity. Activity of each pore can be tracked over time. Figure 1B shows sample time traces from 4 individual Aβ42 pores, highlighting the large variability in their gating properties. We use a large number of such traces to at each recording time to model kinetics of Aβ42 pores, their evolution, and voltage dependence (Table 1 and 2).

**Table 1.** Time evolution of Aβ42 pores function. Time at which the activity of Aβ42 pores were recorded after cell’s exposure to Aβ42 oligomers (row 1) and total number of pores observed (row 2) at that time.

|  |  |  |  |  |  |
| --- | --- | --- | --- | --- | --- |
| Time (min) | 25 | 30 | 35 | 40 | 50 |
| # of pores | 614 | 237 | 262 | 264 | 383 |

**Table 2.** Voltage dependence of observed number of Aβ42 pores. Voltage at which the activity of Aβ42 pores was recorded (row 1) and total number of pores observed (row 2) at that voltage.

|  |  |  |  |
| --- | --- | --- | --- |
| Voltage (mV) | -20 | -50 | -80 |
| # of pores | 1377 | 4297 | 5438 |

### Idealization of time traces

Modeling the gating kinetics of ion channels requires idealized traces representing SCL in which the channel is gating as a function of time. To idealize the raw time traces of several thousands of Aβ42 pores produced by CellSpecks, we used another in-house java-based software called TraceSpecks [54]. TraceSpecks uses maximum likelihood formalism for separating signal from a noisy raw time trace with drifting background. It can automatically idealize thousands of raw traces on the fly. After removing experimental biases and noise (Figure 1B, green) from the raw time trace (Figure 1B, blue), an idealized trace (Figure 1C) revealing various SCLs in which the pore is gating is generated. A zoomed-in version of the idealized trace clearly shows that the pore is gating in up to 5 SCLs (Figure 1D). TraceSpecks also saves the dwell times in different conducting states and P_o_ for each pore, which can be later used to develop single-pore models.

A summary of the analysis given by TraceSpecks is presented in Table 3 and 4. For example, from the movie stack recorded 25 minutes after exposing the cell to Aβ42, a total of 614 pores were analyzed (Table 3). Out of 614, there are 359 pores of Type 1 (pores that open to a maximum SCL of 1), 137 pores of type 2 (pores that open to a maximum SCL of 2) and so on. Overall, this is the equivalent to 195.80 minutes of single channel data obtained using conventional electrical patch clamp technique. Similarly, for evaluating voltage dependence of Aβ42 pores activity, we analyzed data on a total of 11932 active pores at -20, -50 and -80 mV (Table 4). This is the equivalent of 3960 minutes of single channel data. It is interesting to note that for Aβ42, not only the total number of pores increases with time (Table 3, column 2), the numbers of different types pores individually also keep on increasing with time (Table 3, columns 3-7). For voltage dependent experiments, the total number of active Aβ42 pores increases as we increase voltage towards more negative values (Table 4, column 2). Note that the oocytes used for the modeling here were selected from our previous study where we characterized the formation and activity of Aβ pores performing many experiments on over 30 oocytes [47, 48]. Aβ pores in all experiments revealed consistent gating kinetics and the data used for modeling is carefully selected from all those experiments.

**Table 3.** Time evolution of the activity of Aβ42 pores. Column 1: Time at which the activity of Aβ42 pores was recorded from an area of 42×42 μm^2^ after addition of Aβ42 oligomers solution. Column 2: Total number of pores observed at each time. Columns 3-7: Number of pores, out of all pores, with maximum SCL indicated in the top row. Column 8: Total time obtained by combining time traces of all active pores at given time.

| Time (min) | # of pores | SCL 1 | SCL 2 | SCL 3 | SCL 4 | SCL 5 | Trace (minutes) |
| --- | --- | --- | --- | --- | --- | --- | --- |
| 25 | 614 | 359 | 137 | 76 | 19 | 23 | 195.80 |
| 30 | 237 | 134 | 59 | 32 | 10 | 2 | 30.32 |
| 35 | 262 | 135 | 75 | 33 | 13 | 6 | 32.99 |
| 40 | 264 | 127 | 83 | 46 | 4 | 4 | 31.28 |
| 50 | 383 | 138 | 126 | 68 | 30 | 21 | 42.31 |
| total | 1760 | 893 | 480 | 255 | 76 | 56 | 332.70 |

**Table 4.** Voltage dependence of the activity of Aβ42 pores. Column 1: Applied voltage at which the activity of Aβ42 pores was recorded from an area of 42×42 μm^2^ after the application of Aβ42 oligomers. Column 2: Total number of pores observed at each applied voltage. Columns 3-4: Number of pores, out of all pores, with the maximum SCL indicated in the top row. Column 5: Total time obtained by combining time traces of all active pores at given time.

| Voltage (mV) | # of pores | SCL 1 | SCL 2 | Trace (minutes) |
| --- | --- | --- | --- | --- |
| -20 | 1408 | 1377 | 31 | 465.11 |
| -50 | 4545 | 4297 | 248 | 1509.00 |
| -80 | 5979 | 5438 | 541 | 1986.00 |
| total | 11932 | 11112 | 820 | 3960.11 |

Traces were separated into five groups, for pores having a maximum SCL of up to 1, 2, 3, 4, and 5. For example, the trace shown in Figure 1D was included in the group of pores with a maximum SCL of 5 (type 5). For the rest of the paper, pores with SCL 1, SCL 2, SCL 3, SCL 4, and SCL 5 are referred to as type 1, 2, 3, 4, and 5 respectively.

### Model selection

We employed the maximum likelihood-base method [50, 51], implemented in the open-source software QUB [55], to fit a Markov chain model to the idealized traces representing the SCL in which a pore is gating as a function of time. From the idealized data of all pores at a given time, we found that the transitions are predominantly between the adjacent SCLs with only few direct transitions into states separated by two SCLs (for example SCL 1 to/from SCL 3). We didn’t find direct transitions between SCLs that are more than two SCLs apart (for example SCL 1 to/from SCL 4). Therefore, while building our model, we only considered the transitions between adjacent SCLs.

As in our experiments, a pore can have a maximum of 5 SCLs in addition to a closed state, we derived a simple model having the minimum number of states and connectivity (Figure 2A) and the best model (Figure 2B) as dictated by Bayesian Information Criterion [62]. Nevetheless, we found that the simplest and best models gave similar results, therefore, we only present results from the simple model from here on. At each recording time, we calculated the transition rates between different states, lifetime distributions, probabilities, mean open times, and mean close times for each SCL. For example, we used the time traces from all 614 pores to get the transition rates between different states at 25 minutes after Aβ42 application (Table 3). Overall, we obtained five values for each transition rate, corresponding to the five time points indicated in Table 3. As discussed later, these multiple values for each transition rate were then fitted with mathematical formulas to get the final model for the gating of Aβ42 pores at any time after exposing the cell to the oligomeric peptide.

**Figure 2.**
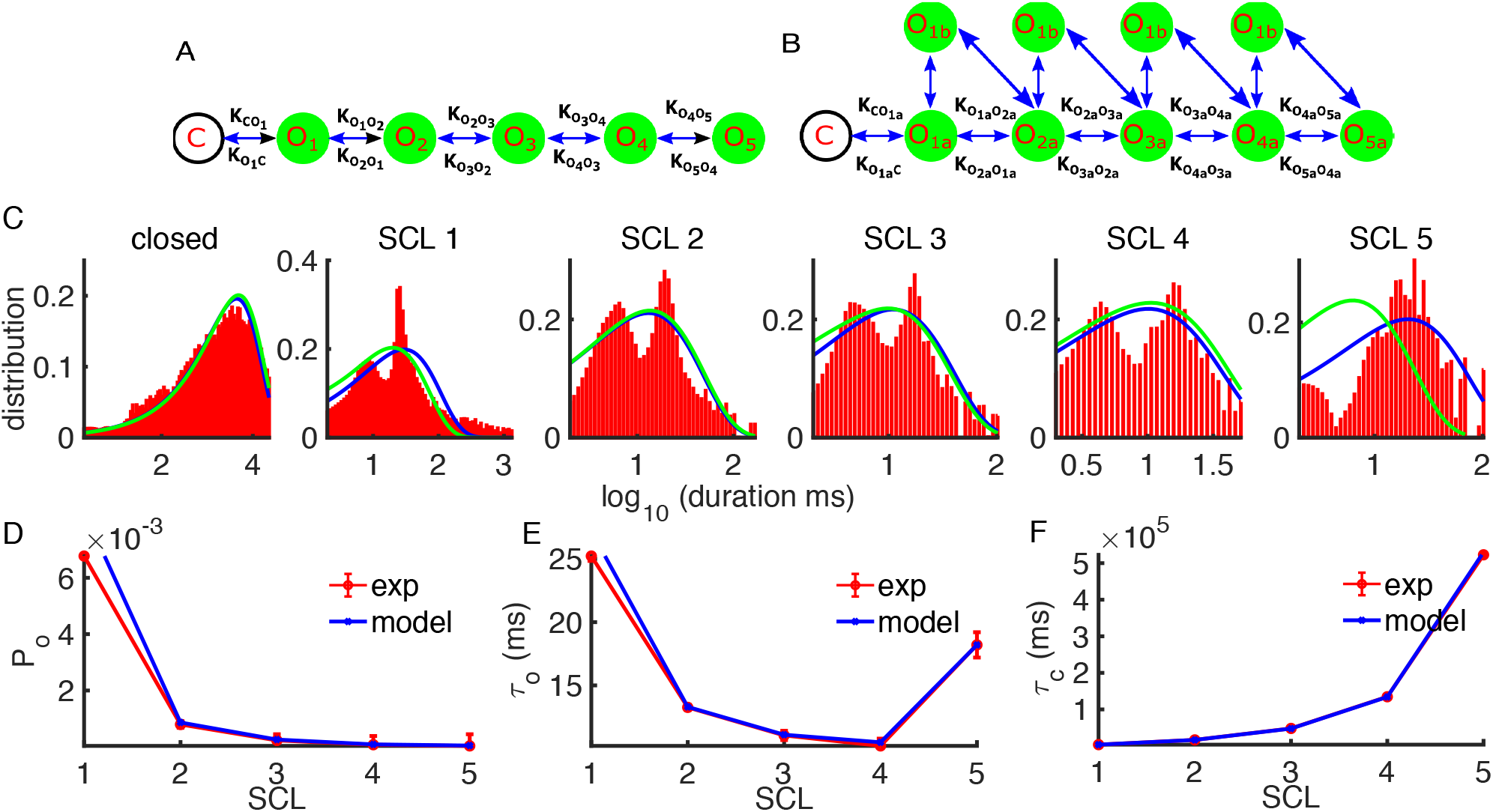
Markov chain model fitting for Aβ42. A) Simple model with one close state and five open states. **B)** Best model according to BIC with extra states added. Rates between newly added states are not shown for clarity but follow similar convention. **C)** Dwell time distributions in closed state and SCL 1, SCL 2, SCL 3, SCL 4 and SCL 5. Red bars are from experiment and blue (green) curves are from the simple (best) model. **D-F)** Average (over all times) statistics of all Aβ42 pores as a function of SCLs: **D)** open probability, **E)** mean open time, and F) mean close time. Red and blue lines are the experimental data and fits from the simple model respectively.

## Results

Results from QUB are shown in Figure 2C, where dwell time distributions for closed state and different SCLs from experiment are shown by red bars and fits from the simplest (blue) and best (green) models are shown by lines. Figure 2D-F show the P_o_, mean open time, and mean close times for different SCLs. As can be seen from Figure 2, the P_o_, mean open times, and mean close time given by the simple model are almost in exact agreement with the experimental data. Although the P_o_ decreases with increasing SCL, we argue that the pores could still have significant Ca^2+^ toxicity while gating in higher SCL (even though for shorter times) due to the fact that the pore conducts more Ca^2+^ in higher SCL. Similarly, mean open time mostly decreases (Figure 2E) and mean close time increases (Figure 2F) as a function of SCL.

In order to check the reliability of our modeling steps, we compared P_o_, mean open time, and mean close time as a function of time as determined by the model with the experimental results as shown in Figure 3A-C. As can be seen, model values agree closely with the experimental results not only for overall P_o_ (Figure 3A), mean open time (Figure 4B), and mean close time (Figure 3C) but also for each SCL (Figures 3D, E) as a function of time.

**Figure 3.**
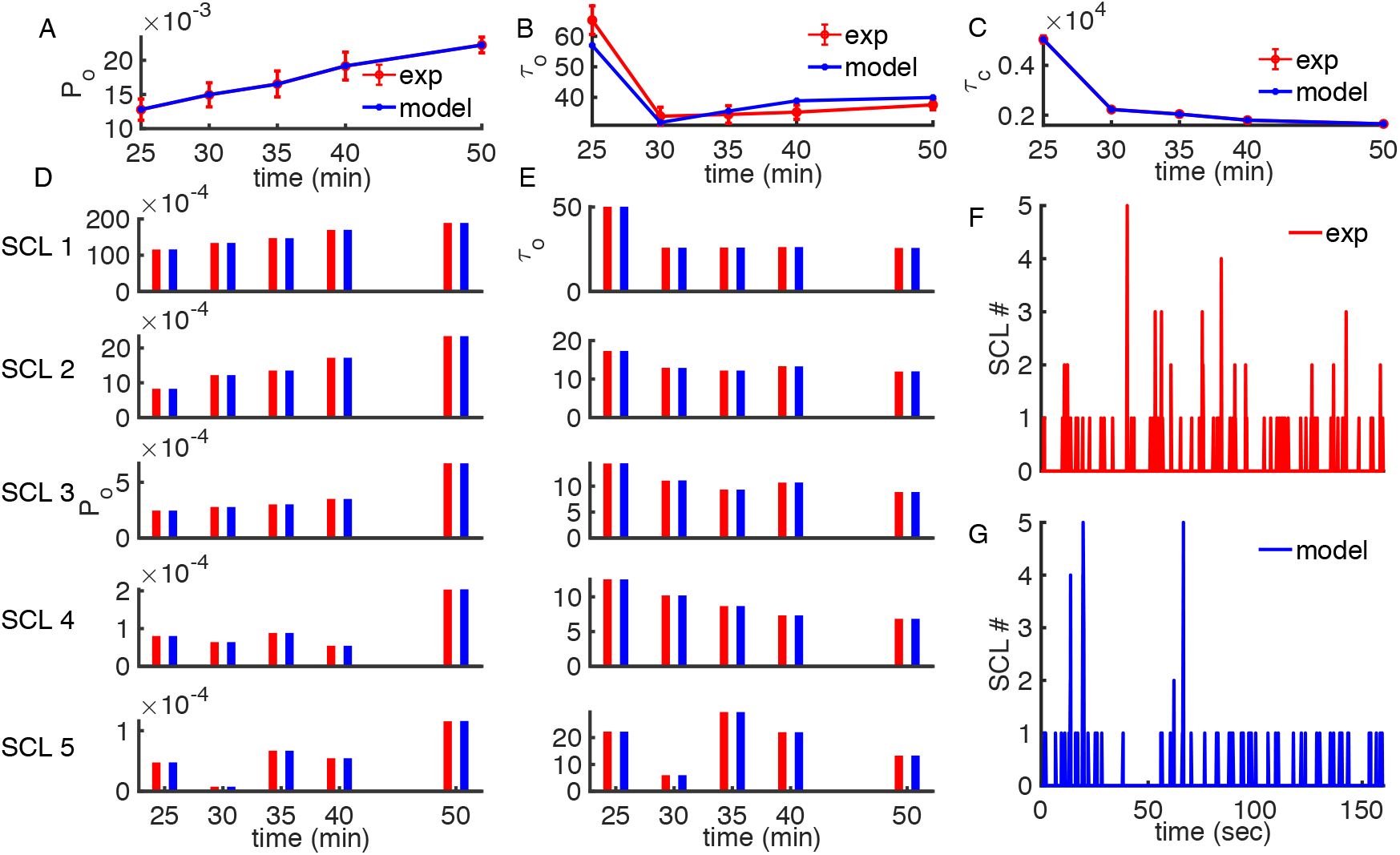
Time dependence of the statistical properties of Aβ42 pores and model time-series traces. Mean open probability **A)**, mean open time **B)** and mean close time **C)** of the pores as functions of time. Mean open probability **D)** and mean open time **E)** for each SCL as functions of time. Example traces from experiments **F)** and from simple model **G)**. In all panels, red lines/bars are from experiment and blue lines are from the simple model.

**Figure 4.**
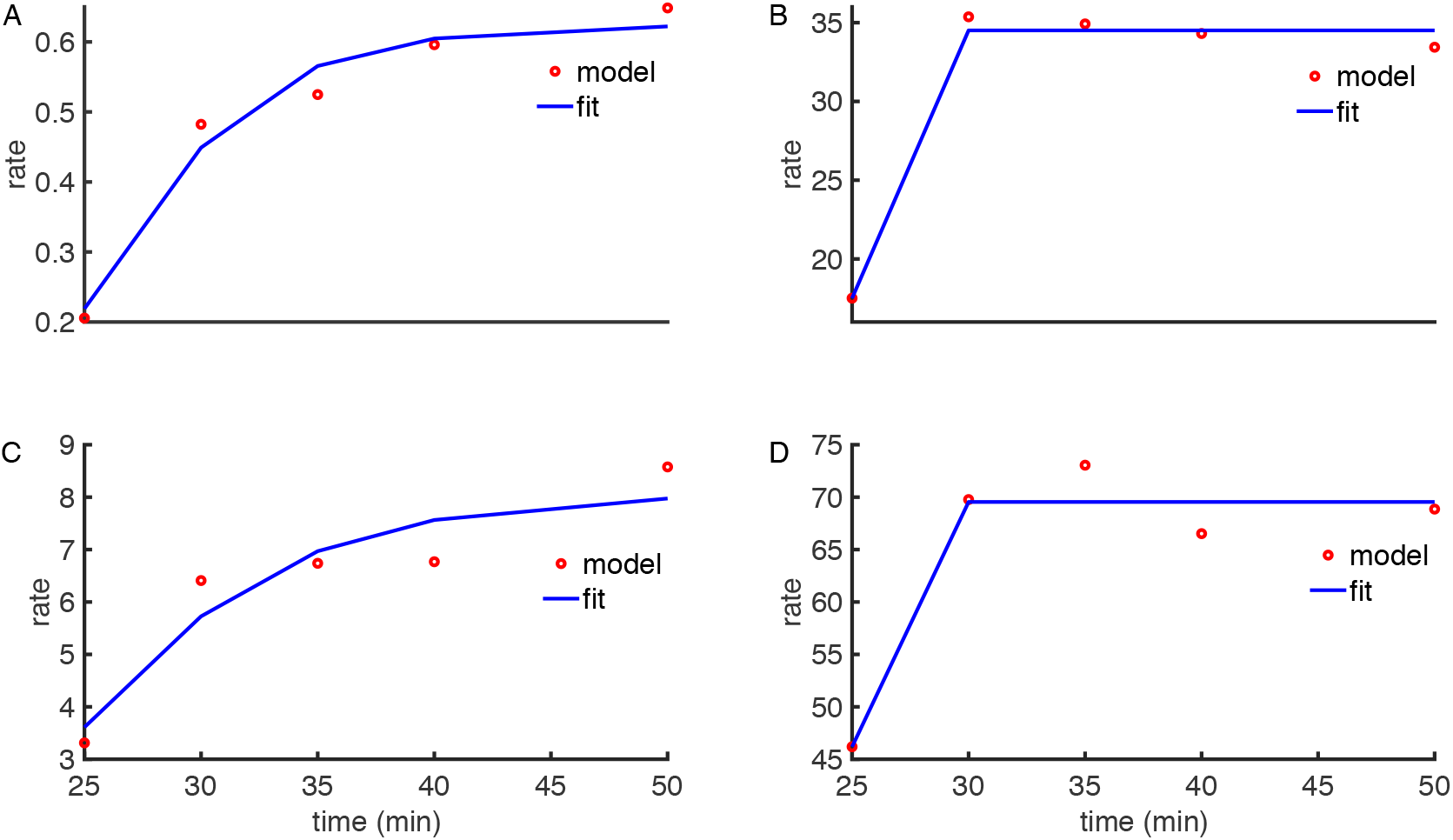
Four parameter Hill’s fits to the transition rates. Transition rates as functions of time after cell’s exposure to Aβ42 oligomers from close state to SCL 1 **A)** and back **B)**. Transition rates from SCL 1 to SCL 2 **C)** and back **D)**. Red symbols are the values given by QUB and blue curves are the fitted values.

### Evolution of Aβ42 pores’ Ca^2+^ toxicity

As pointed out above, QUB gave us multiple values for each transition rate where each value represents the transition probability between two states at a given time. To investigate how the Ca^2+^ toxicity of Aβ42 pores evolve over time, these multiple values (corresponding to multiple movie stacks recorded at different times after cell’s exposure to Aβ42 oligomers) for each transition rate were fitted by four parameter Hill’s function with time as an independent variable. Figure 4A-D show examples of such fittings for transition rates from closed state to SCL1 (and back) and SCL1 to SCL 2 (and back) as functions of time. The resulting Hill’s functions for each transition rate are then used to model the behavior of Aβ pores over longer timescales.

### Simulating single-pore time traces

To test our model further, we next generated time traces representing the gating od a single Aβ42 pore and compared these traces with experimental time traces. We employed the procedure outlined in [63, 64] to simulate these traces. Briefly, if a pore is in state i at time t, the probability with which it makes transition a neighboring state j or remains in state i, within the sufficiently small time interval δt is given by P_ij_ = K_ij_ δt where K_ij_ is the transition rate from state i to j and 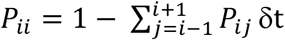. The unit interval is divided into x subintervals of length P_ix_, where x represents the states to which the pore can make transition (including the current state). If a random number drawn from a uniform distribution over the interval δt falls into the subinterval P_ix_, the corresponding transition is performed. The time interval was kept small enough for the linear dependence of P_xx_ on the time interval to remain valid. A sample time-trace for a single Aβ42 pore given by the model is shown Figure 4F (blue). An experimental trace (red) is also shown for comparison, indicating that the trace generated by our model closely resemble to that observed experimentally.

### Estimating the behavior of Aβ42 pores over extended period

We employ the theory of aggregated binary reversible Markov chains to study the dynamical evolution of Aβ42 pores over longer period using occupancy–flux parameterization [50, 56-58]. An Aβ42 pore can gate in 5 open states (5 SCLs) and one closed (C) state as shown in Figure 2A. For simplicity, we represent the closed state by 0 and open states by 1, 2, 3, 4 and 5. In the occupancy-flux parameter formalism, occupancy parameter of a state is given by the product of the forward equilibrium constants for reactions connecting a reference state to that state through an arbitrary path. That is, the un-normalized equilibrium occupancy of a state *i* is given by

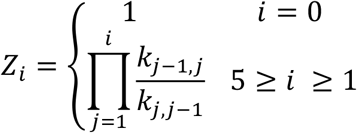

Here *k*_*j* − 1,_*j* (*k*_*j,j*−1_)are the forward/reverse reaction rates from state *j-1* to state *j*. We consider the closed state to be the reference state with un-normalized equilibrium occupancy of 1. The normalized occupancies of any state *i* is given by

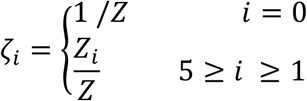

With 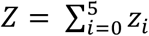

Equilibrium flux between states *i* and *i+1* is given by

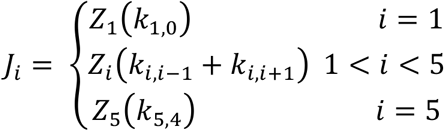

Using occupancy and flux parameters, total P_o_ of a pore is given by

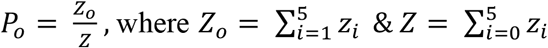

Similarly, P_o_ of a SCL i is given by

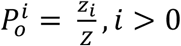

The mean open time and mean close time for a pore can be calculate using the equations

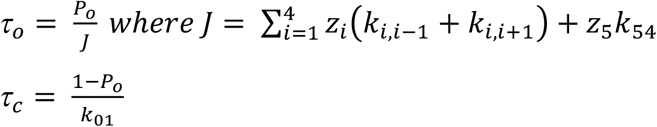

And mean open and close time of a pore in SCL i is given by

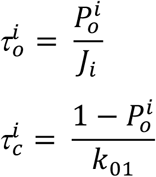

Using the above expression, we computed the statistical properties of Aβ42 pores for up to 500 minutes. These include the total P_o_ (Figure 5A), mean open time (Figure 5B), and mean close time of the pores (Figure 5C). We found that on average the total P_o_ an Aβ42 pore increases with time and reaches an asymptotic value of ∼ 2×10^−2^ (Figure 7A). Similarly, the P_o_ values for each SCL are also shown (Figures 5A) with P_o_ of SCL 1 being the highest and that of SCL 5 being the lowest among all SCLs. As for as the mean open time for each SCL is concerned, we have varying mean open times with SCL 1 having the highest mean open time (23 ms) followed by SCL 2, 5, 3 and 4 as shown in Figure 5B. While the mean open time of the pores as a whole and the mean open times of individual SCLs decrease over time, the rise in P_o_ values mainly results from the steeper drop in the mean close times (Figure 5C). As far as the Ca^2+^ toxicity of these pores is concerned, even though the flux through a pore gating in SCL 5 is five times higher than a pore gating in SCL 1, the pore gating in SCL 1 will be more toxic because the P_o_ of SCL 1 is two orders of magnitude higher than that of SCL5.

**Figure 5.**
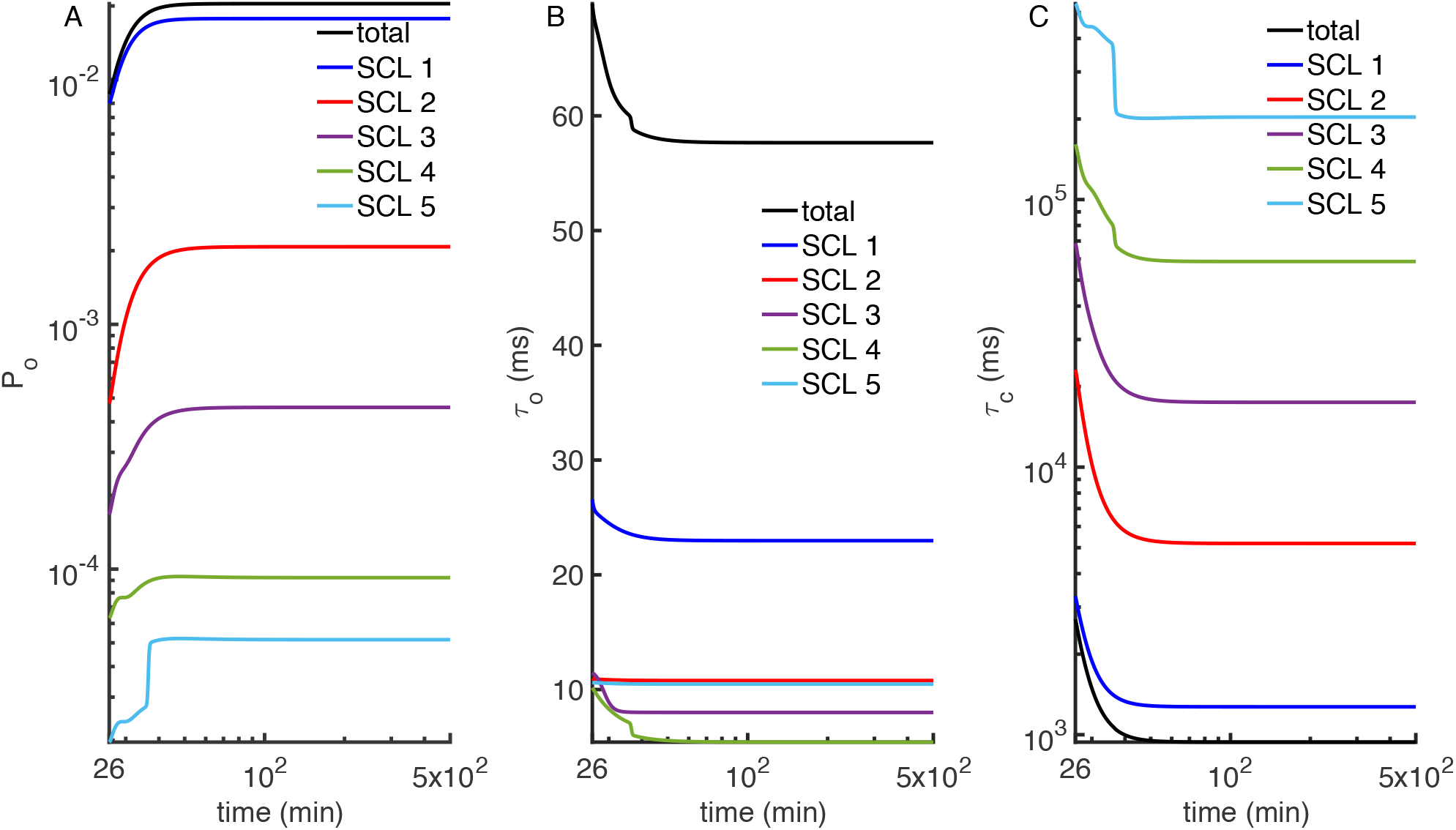
The activity of Aβ42 pores over extended time. **A)** Model prediction for the total P_o_ and P_o_ of each SCL for up to 500 minutes. **B)** Model predictions for the total mean open time and mean open time for each SCL. **C)** Model prediction for the total mean close time and mean close time for each SCL.

### Aβ42 pores cause persistently increasing aberrant cytosolic Ca^2+^ rises

To quantify the changes in cytosolic Ca^2+^ concentration ([Ca^2+^]) due to Aβ42 pores at different times after the cell is exposed to the peptide, we distribute 100 Aβ42 on an area of 100 μm^2^ patch of plasma membrane to mimic experimental pore density. [Ca^2+^] is modeled by reaction diffusion equation given as

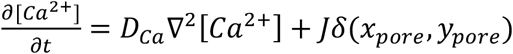

Where *D*_*Ca*_ = 233 *μm*^2^/*sec*, ∇^2^, *δ*, (*x*_*pore*_, *y*_*pore*_) are the diffusion coefficient of Ca^2+^, Laplace operator in rectangular coordinates, Dirac delta function, and the coordinates of a pore. All these parameters are based on experimental observations [65]. Ca^2+^ flux (*J*) through the pore is given by the following equation

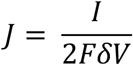

Where *I, F* and *δV* are the current through the pore, Faraday’s constant, and volume of the hemisphere over the pore having a radius of *rpore*. We used *I* = 0.05pA per SCL [48]. That is, the pore allows a current of 0.05pA when gating in SCL 1, 0.1pA when in SCL 2, and so on. *rpore =* 1 nm is based on the value obtained from AFM study of Aβ42 pore [34]. A spatial grid size of 0.05 *μ*m and time step of 1 *μ*sec was used for solving the diffusion equation. The transition rates between different states for individual pores are estimated at each time point are estimated above.

We first quantify the total Ca^2+^ burden due to a single pore when gating in a given SCL by performed solving the above diffusion equation for Ca^2+^ entering through the pore. Figure 6 shows Ca^2+^ rises due to a single opening of a pore while gating in different SCLs. An Aβ42 pore conducting in SCL 1 contributes ∼ 10 μM of Ca^2+^ at the center of pore. As expected, a single opening of the same duration for a pore gating in SCL 5 brings almost five times more Ca^2+^ than that of the pore gating in SCL 1. Similarly, the pore gating in SCL 5 clearly has a wider spatial range of Ca^2+^ rise.

**Figure 6.**
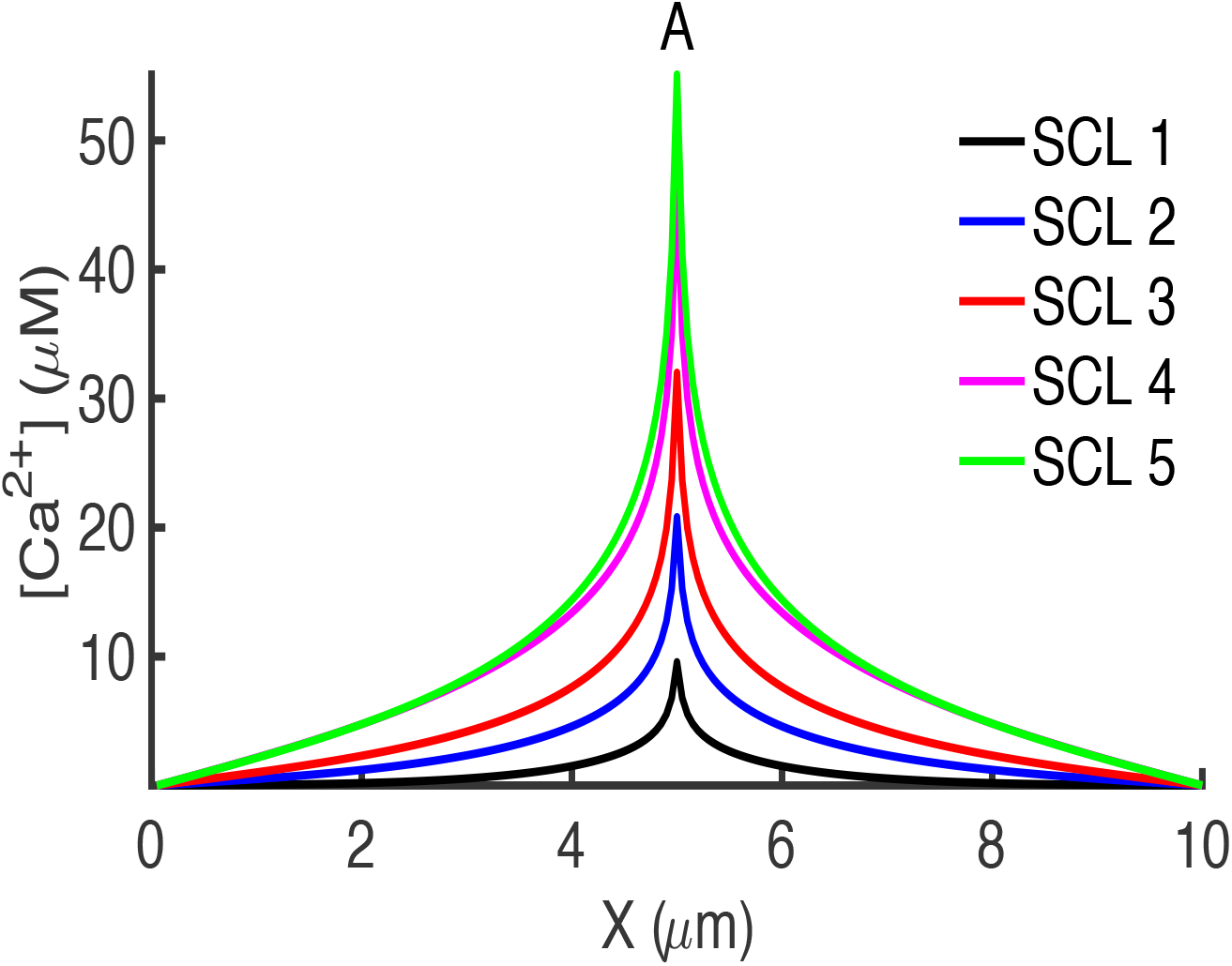
Peak intracellular Ca^2+^ concentration due to Aβ42 pores at the pore location. Ca^2+^ concentration along X-axis passing through the center of the pore due to single opening of Aβ42 pore conducting in SCL 1 (black), 2 (blue), 3 (red), 4 (purple), and 5 (green).

Next, we simulate a 100 μm^2^ patch of plasma membrane with 100 pores (in accordance with experimental pore density of ∼ 1 pore/μm^2^), and generated 10 seconds of data for the overall [Ca^2+^] rise at different times after exposing the cell to Aβ42. Results of these simulations are shown in Figure 7. Figure 7A shows snapshots of the simulated plasma membrane peppered with Aβ42 pores showing the activity of these pores at 40, 45, 50, 55, 60, and 65 minutes after cell’s exposure to Aβ42. There is a persistent increase in Ca^2+^ rise per pore per unit area and the total Ca^2+^ rise due to all pores as the time after cell’s exposure to Aβ42 oligomers increases (Figure 7B, C). [Ca^2+^] increases continuously with the rise of approximately 5 *μ*M due to all pores at 25 minutes to more than 10 *μ*M at 50 minutes after cell’s exposure to Aβ42 (Figure 7B, C). These results clearly indicate that the Ca^2+^ toxicity of Aβ42 pores increases significantly with time.

**Figure 7.**
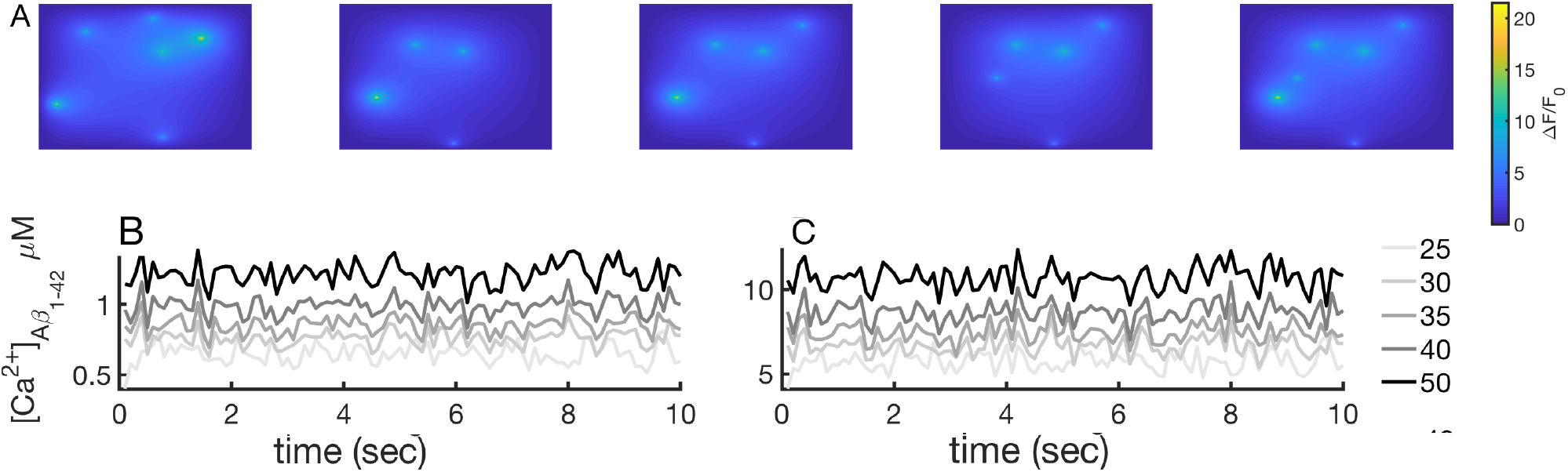
Aβ42 pores leads to a persistently increasing levels of cytosolic Ca^2+^. Activity of 100 Aβ42 pores in a membrane patch of area 10×10 μm^2^ at (left to right) 25, 30, 35, 40 and 50 minutes after cell’s exposure to Aβ42 oligomers. Cytosolic Ca^2+^ accumulation per pore per unit area **D)** and total cytosolic Ca^2+^ accumulated by all 100 pores as a function of time. Each curve represents Ca2+ rises at different time after Aβ42 application as indicated by the legend on the right.

### Dependence of Aβ42 pores’ activity on membrane potential

In order to investigate the effect of driving force on the activity of Aβ42 pores, we recorded the activity of these pores at various voltages as shown in Figure 8 and Table 2. As clear from Table 2, the number of pores detected increases significantly with increasing hyperpolarizing potential. In this set of experiments, we found that almost all pores only go up to SCL 2. Both the P_o_ (Figure 8A, red) and mean open time (Figure 8A, blue) of Aβ42 pores increase, and mean close time (Figure 8A, black) decreases as we make the membrane potential more negative.

**Figure 8.**
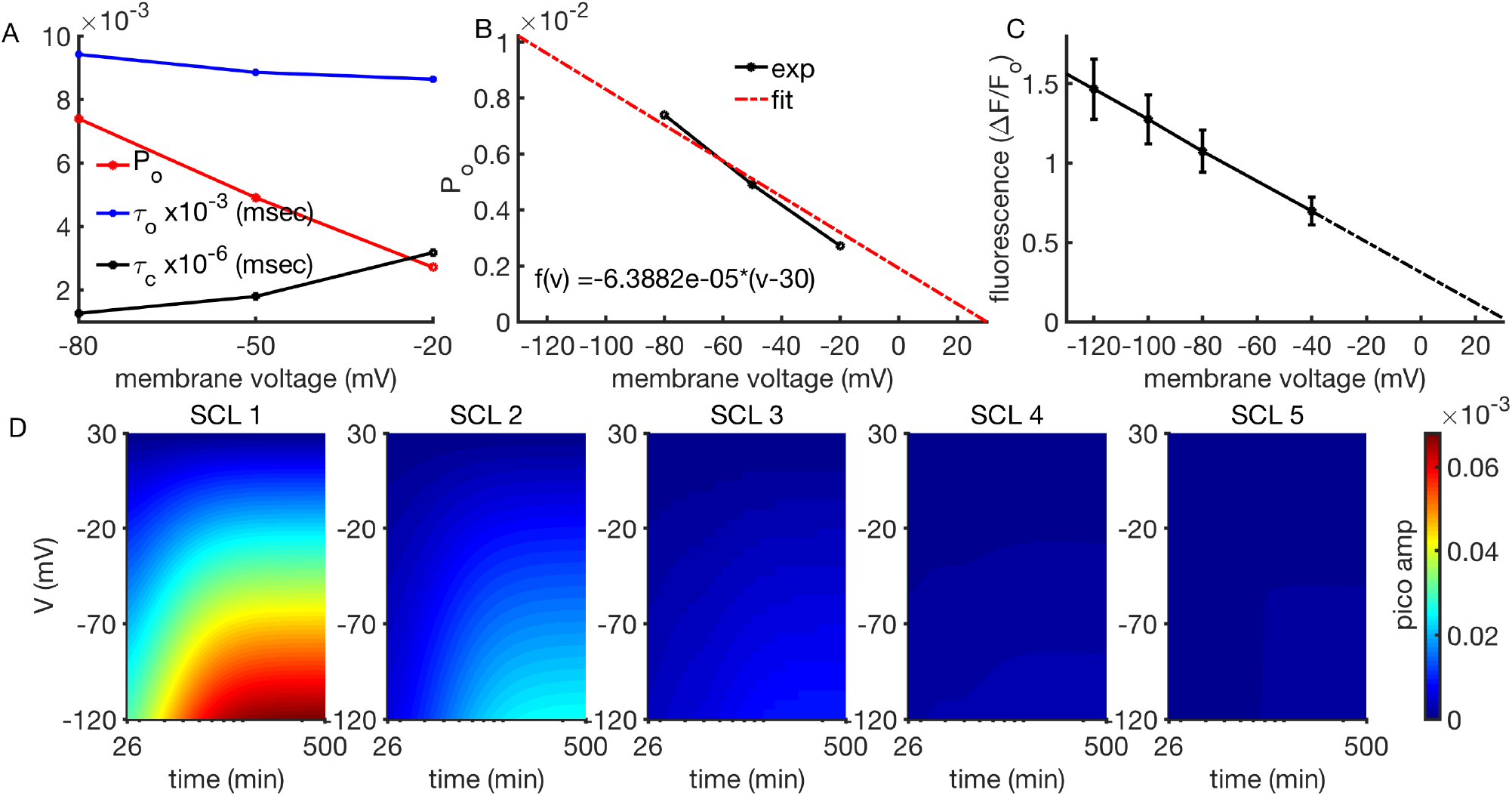
Voltage dependence of Ca^2+^ through Aβ42 pores. **A)** open probability (red line), mean open time (blue line) and mean close time (black line) as functions of applied membrane potential. **B)** Linear fit (red dashed line) to the open probability (black line) with 30 mV reversal potential. **C)** Fluorescence amplitude (ΔF/F_0_) from Aβ42 pores plotted as a function of membrane potential. The fitted regression line extrapolates to 0 at about 30 mV. **D)** Two-parameters voltage-time plots for activity of Aβ42 pores in SCL1, SCL2, SCL 3, SCL 4, and SCL 5 (left to right).

Our previous analysis has shown that the Ca^2+^ flux through Aβ42 pores reverses at membrane potential of 30 mV (Figure 8C) [47, 48]. Thus, we fit the P_o_ (Figure 8B) as a function of membrane potential using linear function of the form:

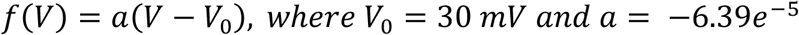

Using Ca^2+^ conductance of 0.4–2 pS for the pores gating in SCL 1 to SCL 5 [48] and reversal potential of 30 mV, we calculated Ca^2+^ current using

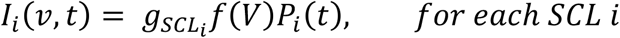

Where 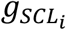 and *P*_*i*_(*t*) are the conductance and P_o_ of the pore in SCL i.

Using *P*_*i*_(*t*) for each SCL from model simulations described earlier, we employed the above function to calculate Ca^2+^current for up to 500 minutes for a range of voltages from -120 mV to 30 mV. Results from these calculations are shown in Figure 8D. The Ca^2+^ flux through Aβ42 pore in each SCL ceases at 30 mV in accordance with the experimental results. Furthermore, for each SCL, Ca^2+^ current through an Aβ42 pore decreases with increasing applied voltage. However, at a fixed hyperpolarizing potential, Ca^2+^ current increases with time and decreases with SCL as shown in Figure 8D (left to right panels).

## Discussion

Growing evidence suggests that abnormal Ca^2+^ signaling due to Aβ pores is detrimental to neuronal functioning and survival. Here, we have shown that PM pores formed by Aβ42 grow in their toxicity with time either due to the formation of pores with higher SCLs in which case they conduct more and more Ca^2+^ or through the growing number of pores resulting in high toxicity. Our robust data-driven modeling approach successfully reproduces the main statistical properties of Aβ42 pores. Namely, mean P_o_, mean open time, and mean close time derived from the model, in general, follow similar trends as experimental observations. Specifically, P_o_ of Aβ42 pores increases with time as predicted by model parameters and observed experimentally as shown in Figure 4. Similarly, pores’ mean close time decreases with time as predicted by model and observed experimentally. Further investigation of the kinetic parameters of these pores at the individual SCLs indicates that the P_o_ for each SCL increases whereas mean open time for each SCL decreases with time after cell’s exposure to Aβ42 oligomers. The persistent increase in the P_o_ signifies the fact that the Ca^2+^ toxicity of these pores increases with time. This finding has significant consequences for the Ca^2+^ toxicity of Aβ42 in Alzheimer’s disease and indicates that while the cell might be able to overcome the aberrant Ca^2+^ rises initially, Aβ42 pores have the potential to produce chronic cell stress over long term, eventually leading to cell’s demise.

After successfully validating experimental observations using our model, we used time dependent Hill’s functions (See Materials and Methods section) in conjunction with occupancy–flux parameter approach to predict the long-term behavior of Aβ42 pores in terms of their level of Ca^2+^ toxicity during disease progression. Our results show that the Ca^2+^ toxicity of Aβ42 pores in terms of the P_o_ continues to grow with time for hours before saturating. Whereas pores with SCLs pores can lead to significant toxicity at longer times, relatively larger number of pores gating in low SCLs are of equal concerns. Furthermore, our quantitative analysis based on Ca^2+^ influx through these pores and diffusion shows that Aβ42 pores can raise local intracellular Ca^2+^ levels by more than two orders of magnitude from resting value of 100 nM to up to 55 μM. Furthermore, our modeling results driven by voltage dependent experiments show that Aβ42 pores are more toxic at hyperpolarizing voltages near resting membrane potentials of neurons.

In summary, we present the first comprehensive data-driven model that not only mimics the gating of Aβ42 pores at a given time but also how their function evolve over long periods after cells exposure to the oligomeric peptide. This model, in conjunction with whole cell model of cell Ca^2+^ homeostasis can provide valuable information on the cytosolic Ca^2+^ disruptions and its downstream effects in Alzheimer’s disease.

